# Filomicelle-Embedded Composite Hydrogels for Localized Gelation within the Anterior Chamber of the Eye

**DOI:** 10.1101/2025.08.19.671150

**Authors:** Hyeohn Kim, Sofia Lara Ochoa, Swagat Sharma, Asmaa A. Youssif, Benjamin R Thomson, Mark Johnson, Evan A. Scott

## Abstract

Nanocarriers hold transformative potential for treating anterior segment eye diseases, yet corneal epithelium impermeability necessitates intraocular injection. Given the discomfort and infection risk, an injectable hydrogel-based depot offers a promising strategy for sustained nanocarrier delivery in intraocular therapy. However, because the aqueous humor is a large, fluid-filled environment, achieving spatially confined gelation remains a key challenge as injected materials rapidly diffuse. Herein, we present a composite hydrogel (C-gel) that enables localized in situ gelation and sustained nanocarrier release within the anterior chamber. This is achieved using poly(ethylene glycol)-b-poly(propylene sulfide) (PEG-b-PPS) filomicelles (FMs), whose filamentous structure confines crosslinking reactions spatially, promoting efficient gel formation. As a result, 90% of the injected polymer is retained within the crosslinked hydrogel matrix. Embedded FMs then undergo oxidation-induced cylinder-to-sphere transitions, facilitating gradual release of micellar nanocarriers. The rheological properties, gelation timing, and microstructure of the C-gel are adjustable, allowing precise control of nanocarrier release dynamics. In vivo evaluation in mice confirmed excellent biocompatibility without inducing intraocular pressure elevation, ocular toxicity, or immune cell infiltration. Sustained release of nanocarriers was observed for over a month under conditions mimicking that of the anterior chamber of the eye, underscoring the potential of C-gels for long-term drug delivery in anterior segment eye diseases therapy.

## 1. Introduction

The anterior segment, comprising the cornea, conjunctiva, iris, ciliary body and lens, plays a crucial role in human vision and ocular health. Anterior segment eye diseases, including cataracts, glaucoma, uveitis, and postoperative inflammation, are the primary causes of vision impairment and account for over 70% of blindness cases worldwide.^[1,2]^ Although numerous topical therapies are approved and available for the treatment of these diseases, many of these drugs exhibit adverse effects on non-target tissues.^[3,4]^ Targeted delivery of these agents using nanoscale drug carriers (i.e. nanocarriers) could emerge as a substantial tool in treatment due to their capacity to minimize side effects in the eye.^[5–7]^ However, the corneal epithelium’s permeability restricts molecules with a molecular weight exceeding 500 Da,^[8,9]^ and nanocarriers cannot traverse this barrier. Consequently, most nanocarriers must be delivered to the anterior chamber via intraocular injection.^[10]^ Considering the discomfort and increased risk of infection associated with frequent intraocular injections, sustained release from intraocular depots over extended periods becomes pivotal for successful pharmacological treatment, yet few options are available for sustained nanocarrier delivery.

Injectable hydrogel platforms offer strong potential for sustained delivery within the anterior chamber.^[11,12]^ However, for clinical translation, there remains a critical need for innovative strategies that enable localized gelation following injection.^[13]^ In conventional hydrogel systems, polymers and crosslinkers are uniformly mixed prior to gelation, leading to global gelation throughout the solution.^[14]^ This process typically requires polymer concentrations above a critical threshold,^[15]^ making it difficult to achieve gelation with only a small amount of polymer. In the context of ocular drug delivery to the anterior chamber of the eye, such a gel must be introduced into the aqueous humor, a saline-like fluid filling this space. Global gelation can cause complications such as visual obstruction and/or blockage of aqueous humor outflow causing elevated intraocular pressure.^[16]^ Therefore, it is important to achieve localized gelation, where a small volume of polymer is injected into a large aqueous environment, and gelation occurs only in the local region without dispersing within the surrounding fluid. To meet this challenge, a platform must enable successful localized gelation under mild physiological conditions, even when a small amount of polymer is injected into a large aqueous phase. Importantly, this must occur without rapid diffusion of the crosslinkers into the surrounding solution, which would otherwise prevent spatially confined gelation.

Poly(ethylene glycol)-b-poly(propylene sulfide) (PEG-*b*-PPS) has demonstrated utility as a novel nanocarrier platform for controlled delivery of therapeutics.^[17,18]^ Stability, high loading efficiency, redox sensitivity, and enhanced endosomal escape of PEG-*b*-PPS nanocarriers have allowed targeted drug delivery and reduction of IOP for anterior segment disease therapy.^[6,7]^ Of note, cylindrical filomicelles (FMs) can reassemble into spherical structures in response to physiological levels of oxidation,^[19,20]^ gradually releasing nanocarriers and enabling their utilization as unique nanocarrier depots for long-term drug delivery. Using this mechanism, PEG-*b*-PPS FM depots have been demonstrated to deliver drugs continuously for months to activate antigen-presenting cells and lymphoid organs such as the spleen and lymph nodes in vivo.^[21]^ Successful in situ gelation and cylinder-to-sphere transitions of subcutaneously administrated FM depots have been thoroughly characterized; however, these gels were designed for crosslinking within dense subcutaneous tissue, and this has limited their translation to aqueous environments such as the anterior chamber of the eye, synovial fluid in joints, and cerebrospinal fluid.

Here we describe a composite hydrogel (C-gel) capable of localized in situ gelation following intracameral injection. This C-gel combines PEG-*b*-PPS filomicelles (FMs) and multi-arm PEG linkers to enable in situ gelation within the aqueous humor via rapid cross-linking and physical entrapment of FMs (Figure 1a). The nanofilamentous architecture of FMs not only promotes spatial confinement of the crosslinking reaction to facilitate localized gelation, but also gradually reassembles from cylinders into spherical micelle nanocarriers, enabling sustained intraocular nanocarrier delivery. The use of DBCO-azide click chemistry fundamentally prevents nonspecific intraocular binding that could arise from less specific chemistries, like vinyl sulfone groups used in previous studies.^[19,21]^ Additionally, we evaluate the rheological properties, microstructure, and gelation dynamics of the C-gels, which can be finely tuned by modifying the ratio of FMs to PEG linkers, total polymer concentration, and temperature to control release kinetics. The potential of C-gels for extended drug delivery in an environment mimicking aqueous humor and its responsiveness to oxidative conditions are assessed. Additionally, in vivo biocompatibility and stability of intracamerally injected C-gels are evaluated by monitoring IOP, ocular integrity, and absence of immune cell infiltration following intracameral injection in mice. This study represents a pioneering platform for the sustained release of nanocarriers in the anterior chamber, expanding the ophthalmic applications of nanocarriers and hydrogels for long-term treatment of ocular diseases.

**Figure 1.**
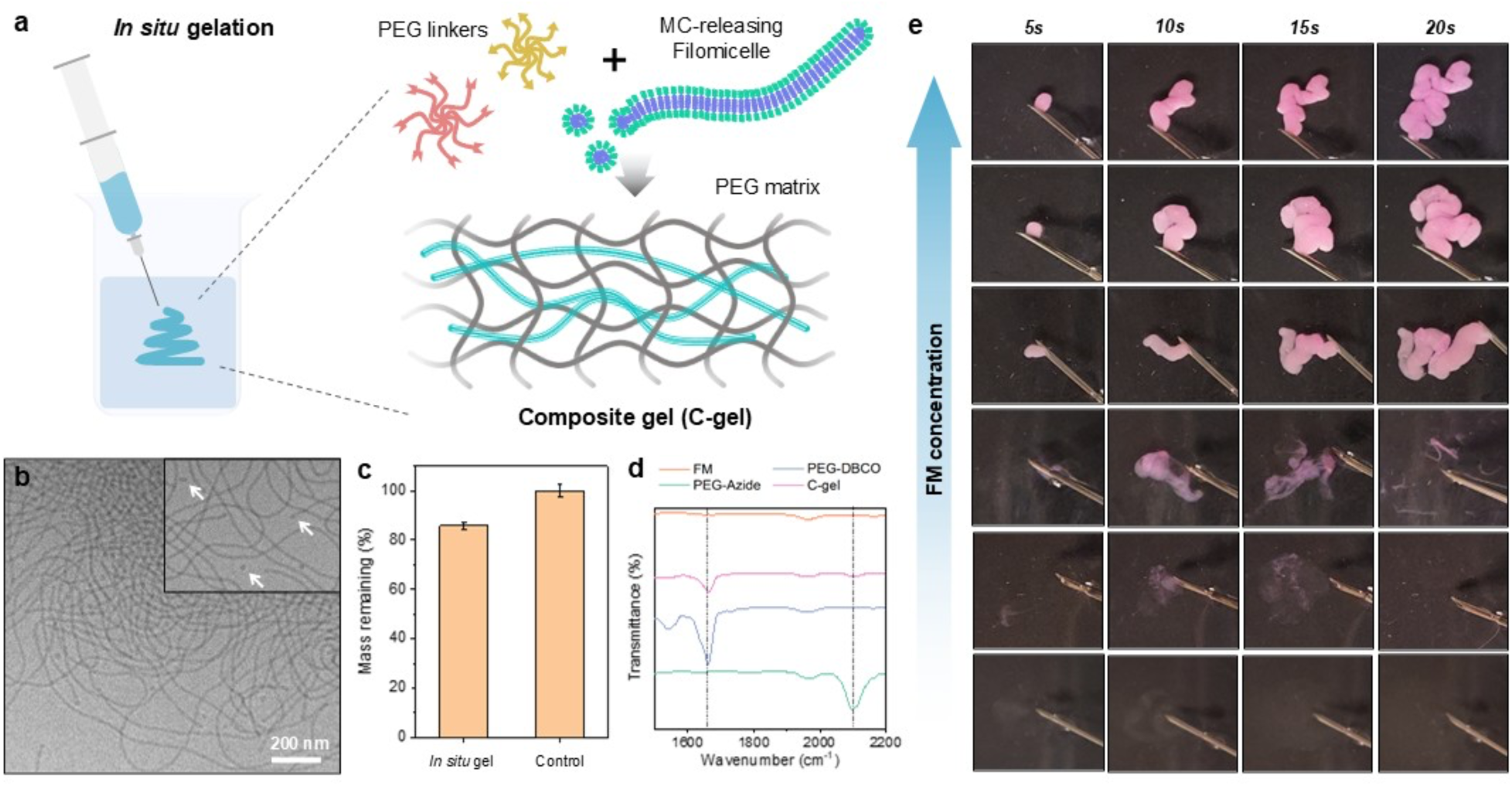
In situ gelation of composite hydrogel (C-Gel) in an aqueous solution. (a) C-gel engineered for in situ gelation within aqueous humor. C-gels are composed of an outer click-chemistry cross-linked PEG matrix with embedded MC-releasing FM. (b) Cryogenic TEM image of the assembled FM structures. Arrows in the enlarged image indicate MCs releasing from the ends of FMs. (c) Mass remaining after in situ gelation of C-gel. Hydrogel was prepared with a FM:PEG linkers ratio of 40:60 (v/v) at a total polymer concentration of 100 mg/mL. Following injection into deionized water, gels were lyophilized after supernatant removal. Controls were lyophilized without removing the supernatant. Error bars represent s.e.m., *n* = 4. (d) Attenuated total reflectance Fourier transform infrared (ATR FT-IR) spectral analysis of the FM (orange line), 8-arm PEG-DBCO (blue line), 8-arm PEG-Azide (green line), and C-gel (pink line). Antisymmetric stretch vibration of the azido group appears at 2,100 cm^−1^. C=C stretching vibration of the aromatic ring of DBCO groups occurs at 1,660 cm^-1^. (e) Effect of FM concentration on in situ gelation of hydrogel in aqueous solution. DiI-loaded FM was prepared via thin film hydration and mixed with multi-arm PEG linker (36 mg/ml) at final FM concentrations of 0, 10, 20, 30, 40, or 50 mg/mL. Mixtures were injected into 37°C PBS at 3 µl s⁻¹.

## 2. Results and Discussion

To create C-gels, a PEG_45_-*b*-PPS_45_ block copolymer (BCP) was synthesized using previous methods with chain length characterized by 1H NMR.^[19,22]^ This BCP is self-assembled into a high aspect ratio filamentous structure through the thin film hydration method, exhibiting micrometer-scale lengths and a core diameter of approximately 20 nm (Figure 1b). Under oxidative conditions, the FMs undergo a thermodynamically driven transition from cylinders to spheres due to alterations in surface tension resulting from the conversion of propylene sulfide to more hydrophilic sulfoxide and sulfone derivatives.^[19]^ This gradual release of spherical micelles (MCs) from the FMs facilitates their long-term therapeutic applications in composite gels. The release of MCs from the tips of assembled FMs can be shown in a magnified cryoTEM image (Figure 1b, inset). Sequential mixing of FM with DBCO- and Azide-PEG linker solution, followed by injection into Dulbecco’s phosphate-buffered saline (PBS), results in in situ hydrogel formation (Figure 1a). The hydrogel crosslinking relies on copper-free DBCO-azide click chemistry under physiological conditions. When the linkers and FMs are combined together, the two linkers are rapidly crosslinked, forming a PEG matrix, physically entrapping the FMs within, and transitioning from sol to gel.

In addition to facilitating the release of MC, high aspect ratio FMs play a crucial role in localized gelation within aqueous volumes. In situ gelation of C-gel was evaluated in PBS with pH, viscosity, and density similar to aqueous humor.^[23]^ Specifically, the FM solution was thoroughly mixed with DBCO- and azide-linkers at a 1:1 molar ratio of functional groups. Using a syringe equipped with a 21G needle, the mixture was injected into PBS at 37°C to observe gelation behavior. In situ gelation was assessed across FM concentrations ranging from 0 to 50 mg/ml under a fixed PEG cross-linker concentration of 36 mg/ml. When only the PEG cross-linker was present or when the FM concentration was below 20 mg/ml, the injected mixture diluted into the PBS without forming a solid gel (Figure 1e). By contrast, at FM concentrations of 30 mg/ml or higher, the mixture rapidly gelled into a solid upon injection, forming a worm-like in situ hydrogel.

Notably, when a spherical MC solution is used instead of FMs, the mixture disperses in water rather than forming a solid gel, even at MCs concentrations above 30 mg/mL (Figure S1). It is suggested that the anisotropic morphology of FM enhances friction and interlocking, possibly due to filament reptation, leading to increased viscosity, and thereby slowing the injected mixture from dispersing in the solvent to promote local crosslinking of the multi-arm PEG into a solid gel (Figure S2). Following gelation, removal of the supernatant and subsequent evaluation of the remaining mass indicated that about 90% of the total polymer was incorporated into the in situ gel, demonstrating highly efficient incorporation and minimal dissolution in the solution (Figure 1c). Analysis of the dried hydrogel via FTIR revealed a significantly reduced characteristic signal for azide, indicating quantitative conversion of azide to triazole upon reaction with DBCO-PEG linkers (Figure 1d).

To achieve stability and functionality in hydrogels, tailored physical characteristics suitable for therapeutic applications are crucial. An injectable hydrogel platform operating in ocular environments must rely on in situ hydrogel formation inside the eye, emphasizing the importance of precise control over gelation timing.^[24]^ In C-gels, the tunable characteristics of PEG-based hydrogels include control over gelation time, final network structure, and stiffness by specifying three main parameters: the FM:PEG linkers ratio, total polymer concentration, and temperature. The ratio of FM to PEG linkers is varied at a fixed total polymer concentration of 100 mg/ml, comprising both the FM and PEG linkers. When varying the total polymer concentration, the FM:PEG linkers ratio is held constant at 40:60 (v/v). The adjustability of gelation time was evaluated by varying these parameters at pH 7.4, using a pipetting test (Figure 2a). Experimental results demonstrate that gelation time can be manipulated from tens of seconds to several minutes by controlling the FM:PEG linkers ratio, total polymer concentration, and reaction temperature, with gelation time increasing with higher FM content relative to the PEG linkers, lower total polymer concentration, and lower reaction temperature.

**Figure 2.**
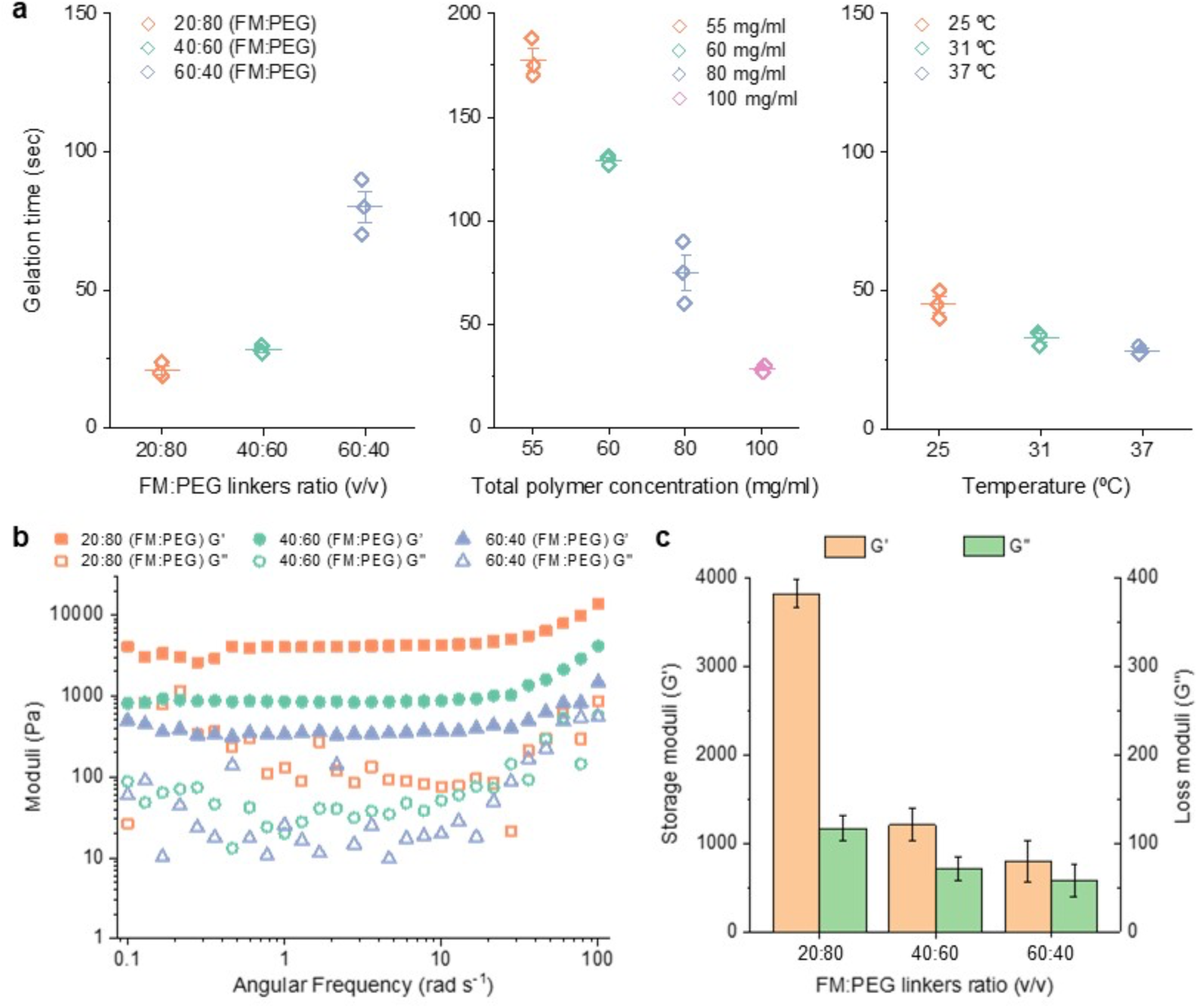
Rheological properties of C-gels. (a) Gelation time of hydrogels determined by a pipetting test. The FM:PEG linkers ratio was varied from 20:80 to 60:40 (v/v) at a fixed total polymer concentration of 100 mg/mL; total polymer concentration was varied from 55 to 100 mg/mL at a fixed FM:PEG linkers ratio of 40:60 (v/v); temperature was varied from 25 to 37 °C. Experiments were conducted in triplicate. The center line and whiskers represent the mean ± s.e.m. (b) Rheological properties of in situ hydrogels with different FM:PEG linkers ratios at 37 °C. Frequency-dependent oscillatory experiments were conducted in a linear viscoelastic regime. (c) Storage moduli (G’) and loss moduli (G’’) of hydrogels with different FM:PEG linkers ratios at 1 Hz (Strain = 0.5%, 37°C). Error bars represent s.e.m., *n* = 3.

The rheological properties of hydrogels can also be easily modulated according to the C-gel composition. For rheological testing, FM, PEG-azide linker, and PEG-DBCO linker solutions were mixed to achieve compositions with FM:PEG linkers ratio from 20:80 to 60:40 (v/v), while maintaining a total polymer concentration of 100 mg/mL. For in situ gelation, hydrogels of varying compositions were injected into PBS at 37°C. After gelation, the hydrogels were collected and evaluated using a rheometer (Figure 2b). The linear viscoelastic region was first confirmed through amplitude sweep, revealing consistent linear viscoelastic regions across all gels (Figure S3). Subsequently, the frequency dependence of the storage moduli (G’) and loss moduli (G’’) of hydrogels in the viscoelastic region was measured at a 0.5% applied strain. For all investigated hydrogel formulations, the storage modulus exceeded the loss modulus across the entire frequency range, indicating the presence of a chemically cross-linked robust fibrous network within the C-gels. The storage modulus (G’) and loss modulus (G’’) of the hydrogels decreased with increasing the FM proportion (Figure 2c). The incorporation of FM is considered to impart both flexiblity and mobility to the hydrogel, rendering the material more pliable. C-gels with all tested FM:PEG linker ratios exhibited superior G’ value compared to previously reported FM-depots,^[19]^ with elasticity comparable to biological tissues.^[25,26]^ Additionally, the G’/G’’ ratios of 40:60 and 60:40 gels, calculated from Figure 2c, ranged from 14 to 17, aligning with characteristics reported for other tissue-mimicking phantoms.^[27,28]^

In C-gels, the pore size and distribution are pivotal design features affecting the diffusion properties of nanocarriers. To evaluate the microstructure of C-gels, we employed cryogenic scanning transmission electron microscopy (cryoSTEM). For this, C-gels with varying FM:PEG linkers ratios were rapidly quenched using liquid ethane on a Cu grid, then loaded into a cryo transfer holder for imaging. The rapid quenching of gel leads to an amorphous solid, disrupting the nucleation of water crystals and allowing for clear observation of the intact hydrogel pore.^[29]^ A controlled warming process at a rate of 3-5 °C/minute under vacuum gradually sublimates the background ice, revealing the hydrogel’s feature. Representative cryoSTEM images clearly visualized the continuous and homogeneous porous structure of the C-gels (Figure 3a-c). Across all gels, sub-micron-sized polygonal pore structures were observed, along with distinct pore-and-wall morphology on the exposed gel surface. Furthermore, depending on the extent of ice sublimation, we noted continuous pore structures within the gel. The sub-micron-sized pores of these gels are considered advantageous for the sustained release of nanocarriers.

**Figure 3.**
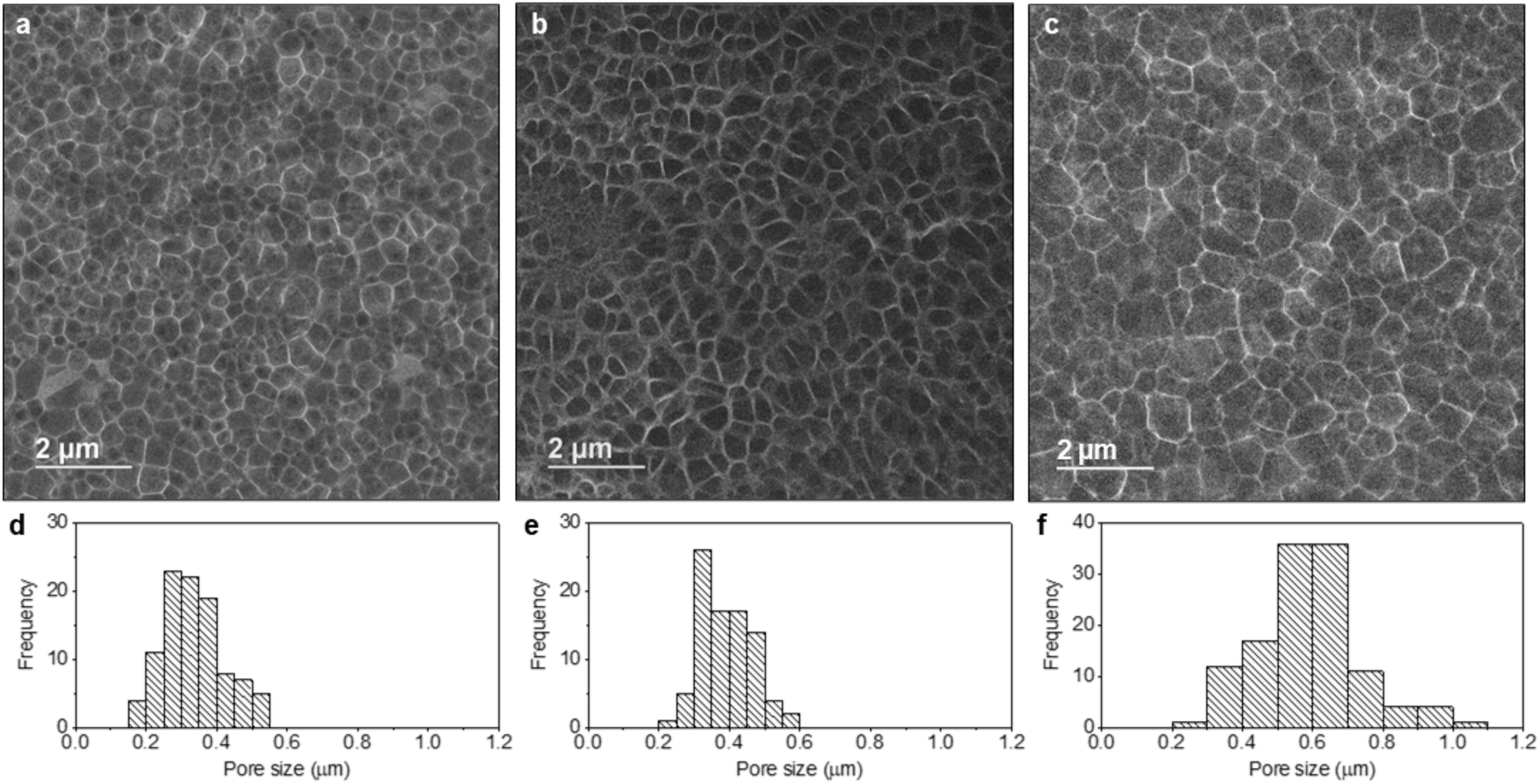
Microstructures of C-gel network. (a-c) Representative cryoSTEM images of *in situ* C-gels prepared at FM:PEG linker volume ratios of 20:80 (a), 40:60 (b), and 60:40 (c) (v/v), with a fixed total polymer concentration of 100 mg/mL. The sub-micro pore structure of the C-gels was revealed through a warming process at a rate of 3-5 °C/minute under vacuum. (d-f) The pore size distributions of C-gels with FM:PEG linkers volume ratios of 20:80 (d), 40:60 (e), and 60:40 (f) were analyzed using ImageJ software.

Pore size ranges and distributions for C-gels with different FM:PEG linkers ratios were analyzed based on cryoSTEM images, followed by further analysis with ImageJ software. An illustration of this approach can be found in the supplementary data (Figure S4). The size analysis showed the increasing tendency of the pore size to increase as the proportion of PEG linkers to FM decreased (Figure 3d-3f). For the 20:80 and 40:60 FM:PEG gels, pore sizes predominantly fell within the range of 0.2 - 0.6 μm, with average pore sizes of 331 nm and 388 nm, respectively. The 60:40 FM:PEG gel exhibited a wider pore size distribution (0.2 - 1.1 μm) with an average pore size of 587 nm. This aligns with previous studies on PEG-based hydrogels, where higher amounts of crosslinkers resulted in smaller pore sizes.^[30–32]^ In addition, the pores were relatively homogenous for both 20:80 and 40:60 gels, as compared with the pores in 60:40 gel, suggesting that the level of homogeneity improved as the proportion of PEG linkers increased. These findings indicate that C-gels not only possess sub-micron pores but also that the pore size distribution and network morphology can be controlled through adjustment of the gel composition, suggesting the potential for a rationally designed hydrogel matrix for sustained nanocarrier delivery.

To assess the ability of C-gels to release MC under oxidative conditions in vitro, we utilized our previously published photo-oxidization method.^[19]^ In this setup, the photo-oxidizer ethyl eosin was loaded into FM to chemically simulate light-induced ROS generation^[33]^ and hasten the oxidation process upon exposure to white light, thereby triggering rapid MC release (Figure 4a). The hydrophobic nature of ethyl eosin enabled localized oxidation and rapid partitioning within the PPS core.^[34]^ C-gels with FM:PEG linkers volume ratios from 20:80 to 60:40, containing 0.75% ethyl eosin by mass were injected into a PBS solution at 37°C for in situ gelation. Following a 30-minute cure, the gel underwent white light exposure at 3.5 mW cm^−2^ for varying durations at room temperature. Subsequent cryoTEM imaging of the supernatant surrounding the C-gels after 24 h irradiation revealed spherical, monodisperse MC release across different scaffold ratios. The estimated average diameters, analyzed using ImageJ, indicated consistent MC release within a size range of 19-23 nm for all gel formulations (Figure 4b-4d).

**Figure 4.**
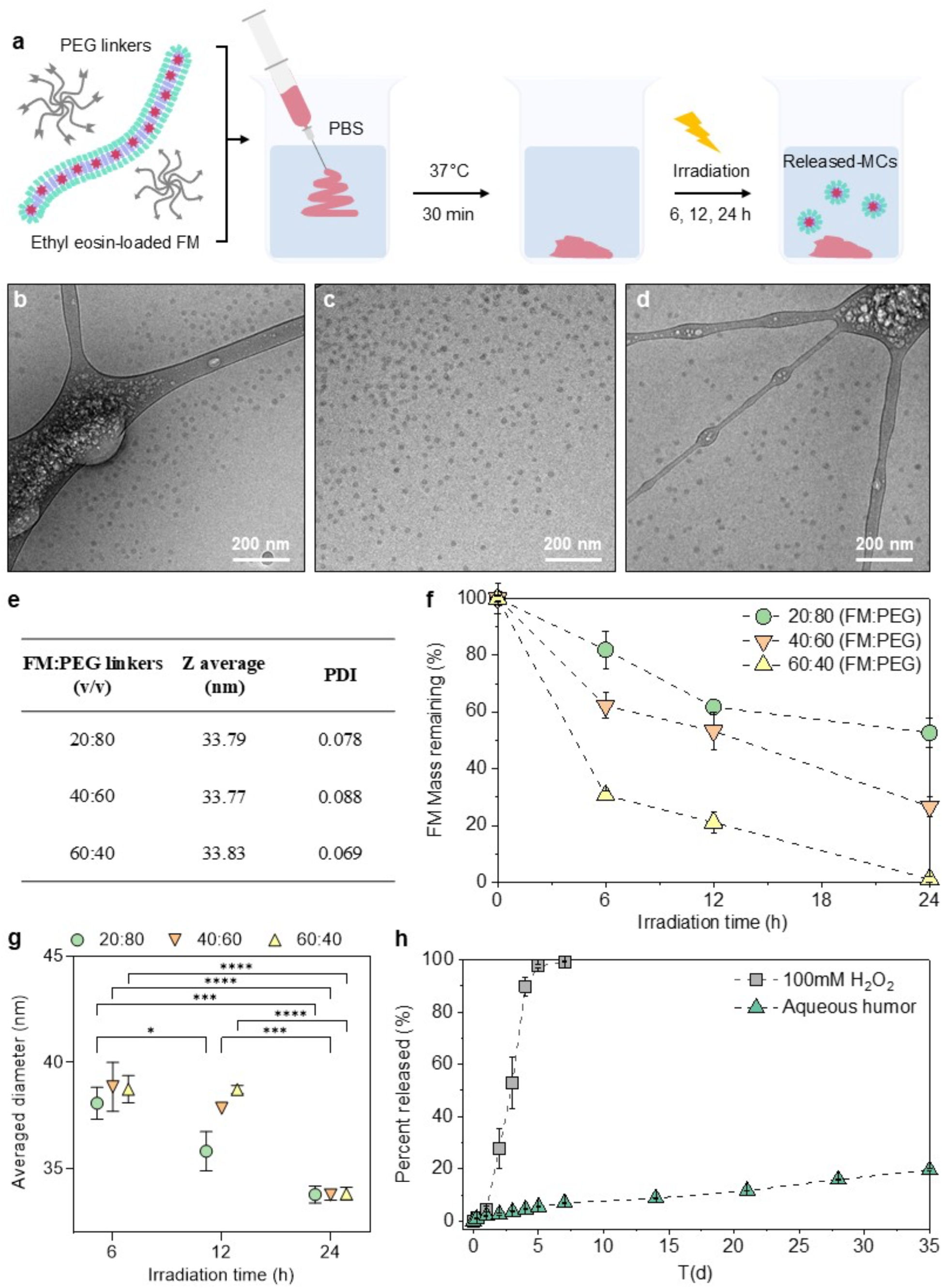
Sustained MC release of C-gels. (a) Schematic of in vitro light-induced degradation of ethyl eosin-loaded FM-embedded hydrogel. (b,c,d) Representative cryoTEM images of released MCs in the supernatant after photodegradation of hydrogels with FM:PEG linkers ratios of 20:80 (b), 40:60 (c), and 60:40 (d) (v/v), prepared at a total polymer concentration of 100 mg/mL. Images were taken on a 200-mesh lacey carbon grid. (e) Averaged diameter and PDIs of MCs released from various C-gels irradiated for 24 h measured via DLS, *n* = 3. (f) Percent mass remaining of FM in hydrogels with FM:PEG linkers volume ratios of 20:80 (green), 40:60 (orange), and 60:40 (yellow) after 0, 6, 12, and 24 h of light irradiation, *n* = 3. (g) Diameters of MCs released from various hydrogels irradiated for 6, 12, and 24 h measured via DLS, error bars represent s.e.m., *n* = 3. Two-way ANOVA was used for statistical analysis. *p* < 0.05 (*), *p* < 0.001 (***), *p* < 0.0001 (****). (h) *In vitro* release kinetics of MCs from C-gels with FM:PEG linkers ratio of 40:60 (v/v) over a 5-week period in rabbit eye aqueous humor and 100 mM H₂O₂ solution. Depicted are average values and error bars represent s.e.m., *n* = 3.

The size distribution of the released MCs was further confirmed by dynamic light scattering (DLS) measurements. The consistent nanoparticle size of approximately 33 nm, regardless of the C-gel composition, supports that the released MCs originated from FMs within the hydrogel (Figure 4e). It is noteworthy that the polydispersity index (PDI) of the released MCs was below 0.1, indicating high uniformity, a notable characteristic of MCs formed via cylinder-to-sphere transitions.^[19]^ The diameters measured by DLS are larger than those measured by cryoTEM. This discrepancy between the two techniques may be due to the bias of DLS measurements for larger higher scattering structures as well as the low contrast of the PEG corona in cryoTEM micrographs,^[35]^ which leads the ImageJ analysis to measure the size of the hydrophobic PPS core of MCs. Furthermore, cryoTEM images sample only a small fraction of the solution, while DLS measurements represent the entire volume.

To examine the impact of the composition of C-gels on delayed release kinetics, we determined the mass of FM remaining within the hydrogel after 6, 12, and 24 h of light exposure (Figure 4f). Notably, altering the FM:PEG linkers ratio from 20:80 to 60:40 (v/v) resulted in an approximately 50% increase in FM degradation rates. These variations in release rates, as illustrated in Figure 3, are attributed to the changes in the relative amount of crosslinkers, affecting the hydrogel’s pore size. Furthermore, extended light exposure beyond 72 h revealed complete MC release without residual FM across all gel formulations (Figure S5). The average hydrodynamic size of MC released over time under light irradiation was analyzed by DLS measurement in all gel formulations (Figure 4g). After 6 h of light exposure, released MC showed an average size ranging from 34 to 38 nm. As irradiation time increased, the size decreased, with an average size of 34 nm observed after 24 h. This size reduction between 6 h and 24 h was statistically significant across all formulations. The formation of a thicker hydrophilic corona due to increased oxidation is consistent with previous findings, which showed that such changes induce the generation of smaller MCs with higher curvature.^[21]^ Together, these findings underscore the ability of C-gels to finely regulate MC release dynamics in oxidizing environments by modulating gel compositions, highlighting its potential for sustained monodisperse nanocarrier delivery.

Furthermore, we demonstrated the sustained release properties of C-gels for continuous nanocarrier delivery in rabbit eye aqueous humor. C-gel with FM:PEG linkers ratio of 40:60 (v/v) underwent gelation upon injection into rabbit aqueous humor solution and 100 mM H₂O₂ solution, with subsequent assessment of MC release over a 5-week period (Figure 4h). The H₂O₂ solution served as a model for reactive oxygen species levels,^[34]^ where high H_2_O_2_ concentrations expedited FM oxidation, thereby accelerating MC release. As a result, complete MC release was observed within one week in 100 mM H₂O₂. However, in aqueous humor containing antioxidants such as glutathione and ascorbic acid, where ROS levels remain in the μM concentration range,^[36]^ the release rate significantly declined, with only 20% released over a 5-week period, indicating remarkable stability. Together, these findings underscore the potential of C-gel as a controlled delivery system capable of sustained release of drug-loaded MCs in the anterior chamber for over a month.

Given the possibility of intraocular hydrogels to adversely affect aqueous humor dynamics, we investigated the IOP response following intracameral injection of C-gel in mice. Volumes of 0.3 μL and 0.6 μL of C-gel with a FM:PEG linkers ratio of 40:60 (v/v) were injected into the anterior chamber of mouse eyes, while an equivalent volume of buffered saline solution was injected into the contralateral eyes (Figure S6a). IOP was monitored for 7 days following the baseline measurement. Considering the total anterior chamber volume in mice is approximately 2.4 μL,^[37]^ these volumes correspond to roughly 12.5 - 25% of the total anterior chamber volume. Although hydrogels are often used in studies to create high IOP animal models due to their potential to cause catastrophic elevation of IOP,^[38–40]^ our study observed no such elevation with C-gel (Figure S6b). This suggests that the rapid gelation kinetics of the C-gel prevent it from flowing into the iridocorneal angles prior to solidification, thus avoiding interference with aqueous humor outflow. Specifically, no significant differences in IOP were observed at any time point between the groups injected with 0.3 μL and 0.6 μL and the control group injected with buffered saline solution. The temporary decrease in IOP on day 1 was likely due to the leakage at the injection site, which was resolved by day 2.

The in vivo compatibility of C-gels was investigated through non-invasive imaging techniques and subsequent histological analysis. A volume of 0.6 μL of C-gel with a FM:PEG linker ratio of 40:60 (v/v) was injected into mouse eyes using pulled glass microneedles, while an equal volume of PBS was administered into the contralateral eyes. Seven days post-injection, stereomicroscopy and high-resolution anterior segment optical coherence tomography (AS-OCT) were employed to assess corneal morphology and anterior chamber integrity. Figure 5a and 5c shows representative microscopic images of eyes injected with 0.6 μL of C-gel or saline. In eyes receiving C-gel, an opaque hydrogel was clearly visible within the anterior chamber. The broad presence of C-gel throughout the chamber is attributed to the relatively large injection volume in the context of the chamber’s small anatomical size. Across all eyes, aside from scars induced by needle puncture on the corneal surface, no signs of epithelial defects, cataracts, corneal haze, or pathological angiogenesis were detected. Representative high-resolution OCT images acquired on day 7 (Figure 5b and 5d) confirmed the sustained presence of C-gel within the anterior chamber, underscoring the C-gel’s robustness. Similar to control eyes, those receiving C-gel exhibited normal anterior chamber configuration iris, lens vault, and iris surface. No signs of anterior uveitis or ocular inflammation were observed, with no indications such as aqueous humor turbidity or floating cells present after the injection. Additionally, no signs of corneal thickening, edema, or irregularities in the corneal endothelium were detected, supporting the absence of direct contact between the cornea and the C-gel.

**Figure 5.**
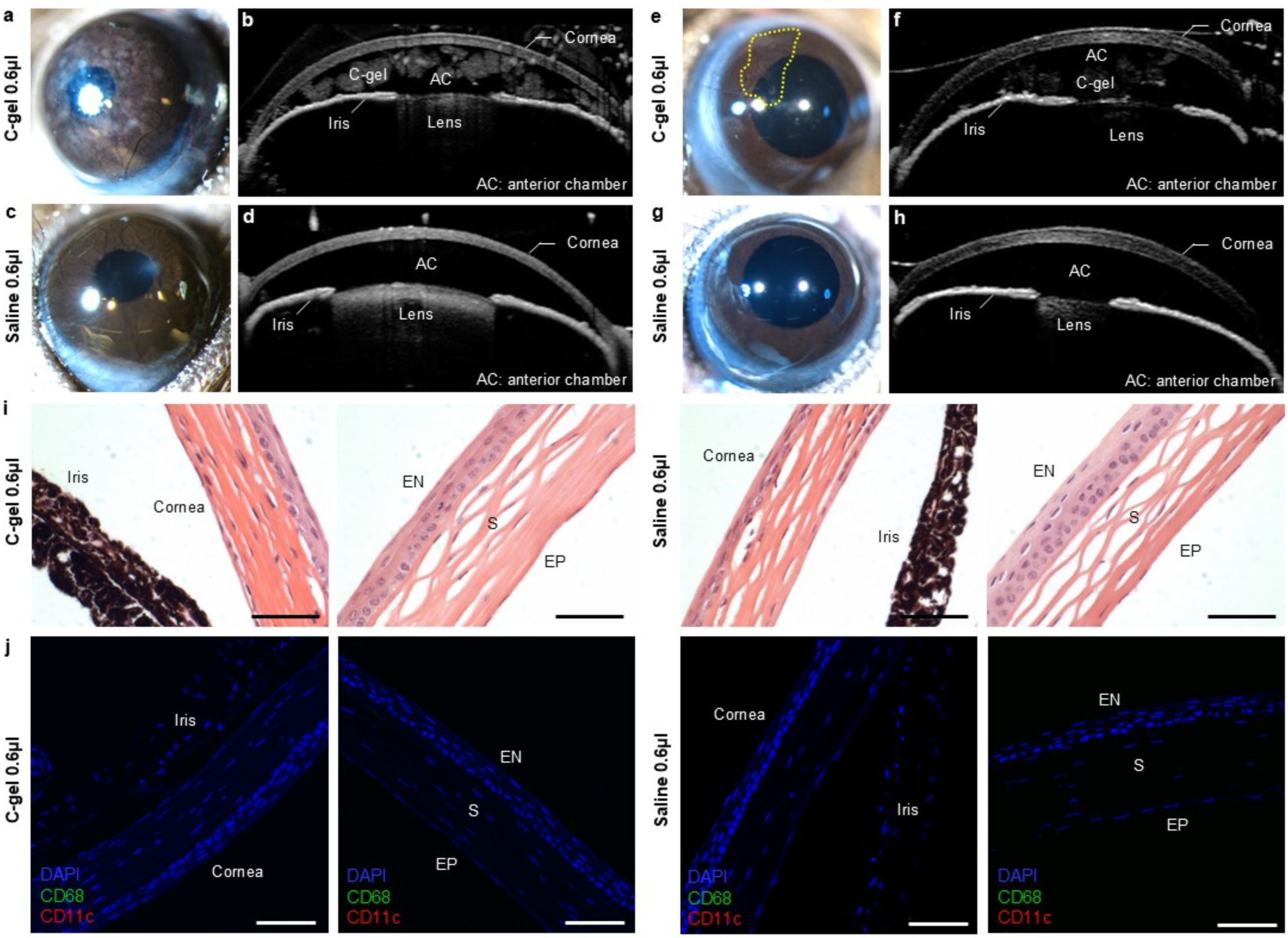
Biocompatibility of intracamerally injected C-gel. (a, c) Optical microscopic images and (b, d) high-resolution anterior segment OCT images of mouse eyes on day 7 after intracameral injection. (a, b) Eye injected with 0.6 µL of C-gel. (c, d) Eye injected with 0.6 µL of saline. Opaque C-gel was clearly visible throughout the anterior chamber. No signs of corneal edema, endothelial disruption, anterior chamber shallowing, iris abnormalities, or aqueous turbidity were detected. (e, g) Optical microscopic images and (f, h) high-resolution anterior segment OCT images of mouse eyes at week 5 post-intracameral injection. C-gel is indicated by yellow dotted lines. (e, f) Eye injected with 0.6 µL of C-gel. (g, h) eye injected with 0.6 µL saline. A decrease in C-gel opacity in the optical image and reduced contrast in the OCT image were observed at week 5. (i) Representative H&E-stained images of the peripheral and central cornea from eyes received C-gel (left) and saline (right). EP, corneal epithelium; S, stroma; EN, corneal endothelium. Scale bar is 50 µm. (j) Representative immunofluorescence images of the peripheral and central cornea from eyes that received C-gel (left) or saline (right). Sections were stained for CD68 (green) and CD11c (red). Scale bar is 10 µm.

To complement the 7-day assessment, imaging at 5 weeks post-injection provided further insight into the ocular biocompatibility of C-gel over time. Stereomicroscopy and AS-OCT revealed no signs of ocular inflammation or structural abnormalities, supporting the continued preservation of anterior segment integrity (Figure 5e-h). Notably, imaging at 5 weeks showed changes suggestive of nanocarrier release. The opacity of the C-gel within the anterior chamber appeared reduced in the stereomicroscopic image (Figure 5e), while the OCT image showed diminished contrast and a more mottled appearance (Figure 5f). Given that OCT contrast reflects refractive index differences associated with density, these changes, together with optical observations, indicate structural alterations in the C-gel, potentially associated with the release of incorporated nanocarriers.

To further investigate potential inflammation and immune cell infiltration induced by C-gel within the eye, histological evaluation was conducted. Eyes that received 0.6 μL of C-gel or saline were harvested 7 days post-injection, followed by tissue sectioning and staining with hematoxylin and eosin (H&E), as well as immunofluorescence labeling for CD68 and CD11c. H&E-stained tissue sections from eyes receiving C-gel showed no apparent signs of structural damage or histopathological abnormalities. Representative sections from the peripheral and central cornea demonstrated a uniformly organized epithelial layer with distinct intercellular borders and appropriate cell density, while the stroma revealed regularly aligned collagen fibers without evidence of edema or disruption (Figure 5i). Immunofluorescence analysis further demonstrated the absence of immune cell infiltration or activation in eyes treated with C-gel (Figure 5j). Maximum intensity projection images of the peripheral and central cornea revealed no visible CD68 or CD11c signals in either C-gel treated or saline treated eyes. No evidence of immune cell clustering or localized accumulation was observed. These histological findings, in conjunction with non-invasive imaging assessments, support the non-immunogenic and non-inflammatory nature of C-gel. Collectively, the results underscore the promising biocompatibility and stability of C-gel for intraocular applications.

## 3. Conclusion

In conclusion, C-gels embedded with PEG-*b*-PPS FMs have been demonstrated as a novel platform for the sustained delivery of nanocarriers within the anterior chamber, enabling the controlled and sustained release of MCs. By leveraging the anisotropic geometry of FMs and click chemistry of PEG linkers, C-gels effectively prevented component dispersion upon injection and enabled spatially confined crosslinking under physiological conditions. CryoSTEM analysis has verified that C-gel possesses sub-micron pores, which is advantageous for the delayed release of nanocarriers. The rheological properties, gelation timing, and microstructure can be intricately designed and adjusted by modulating the composition, polymer concentration, and temperature to enable control over nanocarrier release kinetics. Photo-oxidation tests have demonstrated the hydrogel’s ability to release nanocarriers in response to varying oxidative conditions. Furthermore, C-gels exhibited sustained release of dye-loaded nanocarrier for more than a month in conditions mimicking aqueous humor, thereby showcasing its potential for long-term drug delivery. In vivo studies further supported the biocompatibility, stability, and non-immunogenic nature of C-gel, as evidenced by stable intraocular pressure and the absence of immune cell infiltration on histological analysis.

The therapeutic applicability of this gel could potentially be enhanced by imparting biodegradable properties into the matrix. Introducing photo-cleavable moieties into the multi-arm PEG linkers may allow for photodegradable characteristics through relatively simple chemical modifications. By adjusting the amount of these moieties within PEG linkers, it might also be possible to tune the matrix’s degradation rate, allowing gradual breakdown via ambient visible light or rapid degradation post-nanocarrier release with targeted light exposure. As a pioneering platform for the delayed release of nanocarriers within the anterior chamber, C-gel not only promises to extend drug delivery durations but also to one day enable site-specific anterior eye segment disease therapy within the eye, leveraging the unique advantages of nanocarriers for targeted delivery.

## 4. Experimental Section

### Synthesis of PEG-b-PPS BCP

The synthesis of PEG45-*b*-PPS45 block copolymers was performed as described previously.^[19,22]^ Briefly, methyl ether PEG (MW 2000) was functionalized with the mesylate leaving group, which was then reacted with thioacetic acid to form a protected PEG-thioacetate. Base activation of the thioacetate resulted in the formation of a thiolate anion, which was used as the initiator for ring opening polymerization of propylene sulfide. The reaction was completed with the addition of acetic acid. Degree of polymerization was assessed via H1 NMR (3H methyl ether, 3.36 singlet; 4H PEG―CH2―CH2―, wide peak 3.60–3.64; 1H CH2―CH―CH3 wide peak 2.56–2.65; 2H―CH―CH2―CH―CH3, wide peak 2.82–2.95, 3H―CH2―CH3 wide peak 1.30–1.38).

### FM Assemblies via Thin Film Hydration

FMs were generated via thin-film rehydration. PEG-*b*-PPS was dissolved in ∼2 mL of dichloromethane (5 w/v%) within 2.0 mL clear glass vials. To obtain a homogeneous thin film, a heat gun was used for the removal of dichloromethane. Following the removal of dichloromethane with desiccation, the thin polymer films were hydrated with 1 mL of Dulbecco’s phosphate-buffered saline (PBS) (ThermoFisher Scientific) and gently agitated more than 36 h using a rotator.

### MC Assemblies via Thin Film Hydration

PEG45-b-PPS23 block copolymer was synthesized for the preparation of MC nanocarrier. MCs were assembled using a thin-film hydration method, similar to that used for FM preparation. PEG-*b*-PPS was dissolved in ∼2 mL of dichloromethane (5 w/v%) within 2.0 mL clear glass vials. Following the removal of dichloromethane with desiccation, the thin polymer films were hydrated with 1 mL of Dulbecco’s phosphate-buffered saline (PBS) (ThermoFisher Scientific) and gently agitated more than 24 h using a rotator.

### In situ Gelation of C-gel in PBS

FMs were assembled via thin-film rehydration. For visualization of gelation, lipophilic dye DiIC18(3) (DiI) (ThermoFisher Scientific) was added to the vials for a final DiI concentration, with regard to polymer mass, of 0.067% w/w. DiI loading was confirmed as described previously.^[41]^ For hydrogel formation, the FM solution, 8-arm PEG-Azide (10K, Creative PEGWorks), and 8-arm PEG-DBCO (10K, Creative PEGWorks) were each prepared in PBS and mixed in appropriate ratios to achieve the desired total polymer concentration and FM:PEG linkers ratio. As a typical example, to fabricate a hydrogel with a FM:PEG linkers ratio of 40:60 (v/v) at a total polymer concentration of 100 mg/mL, FM, PEG-azide, and PEG-DBCO solutions were each prepared at 100 mg/mL and mixed in a volume ratio of 20 μL FM, 15 μL PEG-azide, and 15 μL PEG-DBCO solution. The resulting mixture was transferred to a syringe equipped with a needle or a motorized pipette and injected into PBS at 37 °C at a rate of 2 μL/s. The hydrogels were lyophilized and determined by attenuated total reflectance (ATR) FT-IR, and each FT-IR spectrum was recorded with 16 scans in the range of 600–4000 cm^−1^.

### Measurement of Gelation Time

FM, PEG-azide, and PEG-DBCO solutions were mixed in an Eppendorf tube. Gelation time was determined by a pipetting test, which was the time at which the gel could no longer be aspirated with a 10 μL pipette set to aspirate 10 μL of liquid. Temperature control experiments utilized a water bath equipped with a thermocouple. Three replicates of each formulation were performed.

### Viscosity analysis of nanocarrier solutions

Viscosity analysis was conducted at RT in a humidified atmosphere using a modular compact rheometer (Anton Paar MCR302) equipped with an 8 mm parallel plate geometry. FM and MC solutions were prepared via thin film hydration at a final block copolymer concentration of 72.7 mg/mL. 50 µl aliquot of each sample was loaded with a rheometer gap of 0.3 mm. Viscosity was recorded by conducting a shear rate sweep from 0.1 to 100 s⁻¹.

### Rheological Analysis of C-gels

Rheological analysis was conducted at 37 °C in a humidified atmosphere using a modular compact rheometer (Anton Paar MCR302) equipped with an 8 mm parallel plate geometry. FM, azide linker, and DBCO linker solutions were prepared at a concentration of 100 mg/mL and mixed to achieve specific FM:PEG linkers ratios. 55 µl of C-gels with varying FM:PEG linkers ratios, ranging from 20:80 to 60:40 (v/v), were injected into PBS at 37°C for in situ gelation, and collected. The C-gels were evaluated with a rheometer gap of 0.5 mm. An amplitude sweep was performed at a constant angular frequency of 6.28 rad s^−1^ to verify the linear viscoelastic regime. Frequency dependence of the storage and loss moduli was analyzed in oscillatory mode with a 0.5% applied strain. The averaged storage moduli and loss moduli were determined at an angular frequency of 6.28 rad s^−1^, 0.5% strain, 37 °C, *n*=3.

### CryoSTEM Microscopy

For sample preparation, 300-mesh Cu grids with a carbon membrane were glow-discharged for 30 seconds in a Pelco easiGlow glow-discharger at 15 mA with a chamber pressure of 0.24 mBar. The glow-discharge process mitigates the hydrophobicity of the carbon membrane, permitting the aqueous sample to spread evenly over the grid. Approximately 1.5–2 µL of in situ C-gels with different FM:PEG linkers ratios were transferred onto a grid, and plunge-frozen into liquid ethane without blotting using a Vitrobot Mark IV plunge freezing robot. Grids were then loaded into a Gatan 626.5 cryo transfer holder and viewed in a Hitachi HD2300 STEM with a field emission source at 200kV utilizing an SE detector. Prior to gathering image data, the cryo transfer holder was warmed up between −120 °C and −100 °C at a rate of 3-5 °C/minute. This warming process slowly sublimed away background ice under a vacuum to reveal hydrogel features. The pore size of C-gel’s microstructure was evaluated using a manual approach for cryoSTEM images captured at magnifications of 10 000×. A circle drawn using ImageJ software, version 1.54 g, was placed on each pore, which displayed well-defined edges to determine the area which allowed the calculation of the diameter of the pore (details included in Figure S4 in the Supporting Information).

### Photo-oxidation of C-gels

For the photo-oxidation test, ethyl eosin was loaded into FM, and ethyl eosin loading was confirmed as described previously.^[19]^ Visible light was used to generate singlet oxygen to induce FM oxidation and transition to MCs. C-gels including ethyl eosin-loaded FM were incubated in 1 ml of PBS within 2.0 ml clear glass vials, and were exposed to white light (Max-303 Xenon Light Source, Asahi Spectra) at an intensity of 3.5 mW cm^−2^ for 6 to 72 h. Following irradiation, the supernatant was removed and the remaining C-gel was lyophilized. Masses of the lyophilized C-gels were measured to assess degradation, *n* = 3.

### Characterization of Released MCs

Zetasizer Nano (Malvern Instruments) equipped with a 4mW He-Ne 633 laser was used to characterize the size distribution of released MCs in the supernatant of C-gels with different FM:PEG linkers ratios at different time points, *n*=3. The polydispersity index (PDI) was calculated using a two-parameter fit to the DLS correlation data. CryoTEM was also utilized to confirm morphology and size of MCs in the supernatants of C-gels irradiated for 24 h. The average diameter of released MCs from C-gels were analyzed via ImageJ software.

### In vitro Release Study of C-gels

To determine the release kinetics for C-gel, lipophilic dye DiI was loaded into the FM and DiI loading was confirmed as described previously.^[41]^ The FM solution, 8-Arm PEG-Azide (10K, Creative PEGWorks), and 8-Arm PEG-DBCO (10K, Creative PEGWorks) were prepared at a concentration of 100 mg/ml in PBS. For hydrogel formation, 20 µl of FM solution was sequentially mixed with 15 µl of PEG-azide solution and 15 µl of PEG-DBCO solution to achieve the FM:PEG linkers ratio of 40:60 (v/v). 30 µl of the mixture was injected in 0.5 ml rabbit eye aqueous humor solution (Pel-Freez, LLC.) and PBS solution with 100 mM of hydrogen peroxide, and incubated at RT, *n*=3. Aqueous humor solutions were handled in a sterilized laminar flow hood, sealed and retained under sterile conditions. At different time points (from 6 h to 35 days), the samples were gently agitated to ensure homogeneity before collecting 0.25 ml of the supernatant, which was then replaced with 0.25 ml fresh solutions. The amount of DiI-loaded MCs that had been released was then determined by fluorescence measurement (RF-6000 spectrofluorometer) at an excitation of 549 nm and an emission of 565 nm. At different time points (from 6 h to 5 weeks), the samples were gently agitated to ensure homogeneity before collecting 0.25 ml of the supernatant, which was then replaced with 0.25 ml fresh solutions.

### Animals

For this study, 12-week wild type female C57BL/6J mice were purchased from The Jackson Laboratory and were fed a standard diet. The mice were sheltered and maintained in the Center of Comparative Medicine at Northwestern University at 18-23 °C with 40-60% humidity, and 12 h/12 h dark/light cycle. All the experiments were conducted in accordance with animal protocols approved by the Institutional Animal Care and Use Committee (IACUC) at Northwestern University.

### Intracameral Injection

Glass pipettes (Wiretrol II 5 µL, Drummond Scientific) were pulled using a micropipette puller (P-1000, Sutter Instrument) and beveled at a tip diameter of approximately 0.18 mm using a glass grinder (BV-10, Sutter Instrument). The mice were anesthetized with an intraperitoneal injection of ketamine (100 mg/kg) and xylazine (10 mg/kg). The eyes were prepared with 0.5% droplets of proparacaine HCl ophthalmic solution to both eyes prior to injection. The mice were placed on a water circulating heating block during the experiment to maintain body temperature. Pulled glass microneedles filled with C-gel formulation with a FM:PEG linkers ratio of 40:60 (v/v) were connected to a micromanipulator and a pump. The needle was inserted in through the cornea, starting several mm inward from the limbus and run a couple of mm parallel to the corneal stroma before diving down into the anterior chamber. During the injection, the eye was rinsed with Systane eye drops to keep it hydrated. The formulations were administered via perfusion at a flow rate of 500 nl/min and the total volume of injection was 0.3 and 0.6 µl. After the injection, the needles were withdrawn and topical erythromycin antibiotic ointment was applied to both eyes right after the procedure and 24 h later. All the mice were kept on a warm water-circulating blanket until full recovery.

### Intraocular pressure measurements

For IOP studies, 9 12-week wild type female C57BL/6J mice were used. The mice were anesthetized with ketamine (60 mg/kg) and xylazine (6 mg/kg). IOP was measured by rebound tonometry (Tonolab, iCare Inc) at baseline (pre-injection) and time points of 24, 48, 72, 96, day 5, and day 7. Each IOP was the average of 3 measurements, giving a total of 18 rebounds from the same eye per recorded IOP value.

### In vivo biocompatibility analysis

Each mouse received 0.6 µL of C-gel in one eye and PBS in the contralateral eye, following the intracameral injection procedure described above. Briefly, glass pipettes (Wiretrol II 5 µL, Drummond Scientific) were pulled with a micropipette puller (P-1000, Sutter Instrument) and beveled to a ∼0.18 mm tip diameter (BV-10, Sutter Instrument). Pulled glass microneedles filled with either C-gel formulation or PBS were connected to a micromanipulator and a pump, and the formulations were administered via perfusion at a flow rate of 500 nL/min.

After intracameral injection, the eyes of anesthetized mice were thoroughly examined on both week 1 and week 5 (n = 3 per time point) using a stereo microscope (Nikon SMZ800N) equipped with a digital microscope camera (Nikon Digital Sight 10). Anterior segment OCT was also performed using a modular imaging platform (Heidelberg Spectralis HRA+). For imaging, the animals were placed in a custom-built holder which provided precise and reproducible alignment of the animals for OCT imaging. Vertical and horizontal scans, which provide a wider viewing angle to include the entire cornea, were obtained. A minimum threshold of 1.5 times the OCT noise was applied to the images for better clarity and visibility of features.

### Histology and immunofluorescence staining

At week 1 following the intracameral injection of C-gel or PBS, mice (n = 3) were euthanized via carbon dioxide asphyxiation and cervical dislocation. Both eyes were enucleated immediately after euthanasia and subsequently fixed in Davidson’s fixative at RT for 20h, transferred to 4% paraformaldehyde overnight. Following fixation, the tissues were washed with PBS and then dehydrated through a graded ethanol series, embedded in paraffin, sectioned, and stained with haematoxylin and eosin. Stained sections were examined under a light microscope (Olympus BX53).

For immunofluorescence staining, tissue blocks were sectioned at 4 μm thickness and stained with rabbit monoclonal antibodies against CD11c (D1V9Y) and CD68 (E3O7V) (Cell Signaling Technology). Visualization was performed using MACH2 Goat anti-Rabbit HRP polymer (Biocare Medical) and TSA amplification with VIVID650 and VIVID570 (ACD), respectively. Nuclei were counterstained with 4′,6-diamidino-2-phenylindole (DAPI). Antibody specificity was validated using mouse colitis tissue as a positive control, which confirmed robust reactivity (Figure S7). Images were acquired using a Leica SP8 confocal microscope. Z-stacks were captured, and maximum intensity projection images were generated for analysis.

## Acknowledgements

The authors acknowledge a Research to Prevent Blindness unrestricted grant to the Northwestern University Department of Ophthalmology. We also express gratitude to Eric W. Roth for conducting cryoSTEM observations. We acknowledge the assistance of the BioCryo facility of Northwestern University’s NUANCE Center, which has received support from the SHyNE Resource (NSF ECCS-2025633), the IIN and Northwestern’s MRSEC program (NSF DMR-2308691). Comparative histopathology and molecular phenotyping services were provided by the Northwestern University Mouse Histology and Phenotyping Laboratory (MHPL) which is supported by NCI P30-CA060553 awarded to the Robert H Lurie Comprehensive Cancer Center.

## Author Contributions

H.K. and S.L.O. contributed equally to this work. All authors have given approval to the final version of the manuscript.

## Supporting Information

In situ gelation of hydrogel in PBS, Viscosity of FM and MC solutions over a range of shear rates, Strain-dependent oscillatory rheology of C-gels, Examples of the evaluation of pore size, Percent mass remaining of FM in C-gels, IOP response following intracameral injection of C-gel, Positive control for CD68 and CD11c antibody staining. (PDF)

---

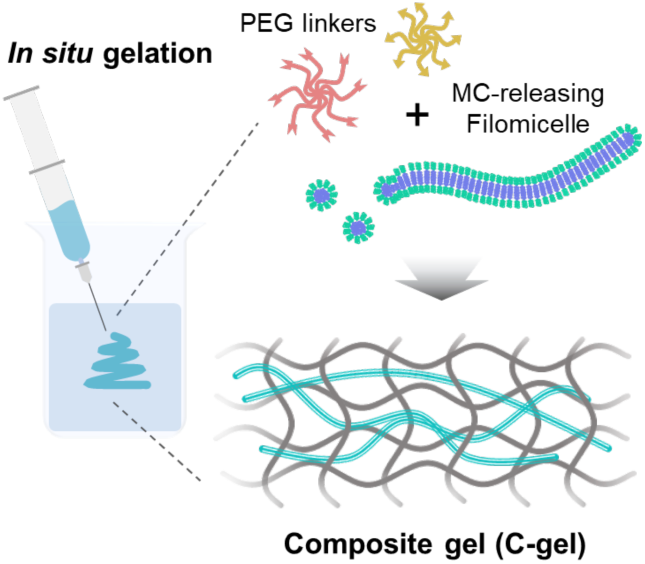
A composite hydrogel (C-gel) incorporating PEG-*b*-PPS filomicelles enables localized in situ gelation and sustained nanocarrier release within the anterior chamber. The filamentous structure promotes spatial confinement of crosslinking even upon injection into large aqueous volumes and gradually transitions into spherical micelles. Designed for intracameral injection, C-gel offers a promising platform for sustained nanocarrier delivery in anterior segment eye disease applications.

