## Supplemental Data for "Filomicelle-Embedded Composite Hydrogels for Localized Gelation within the Anterior Chamber of the Eye"

**CONTENT**


**Figure S1.** In situ gelation of hydrogel in PBS.

**Figure S2.** Viscosity of FM and MC solutions over a range of shear rates.

**Figure S3.** Strain-dependent oscillatory rheology of C-gels.

**Figure S4.** Examples of the evaluation of pore size.

**Figure S5.** Percent mass remaining of FM in C-gels.

**Figure S6.** IOP response following intracameral injection of C-gel.

**Figure S7.** Positive control for CD68 and CD11c antibody staining.

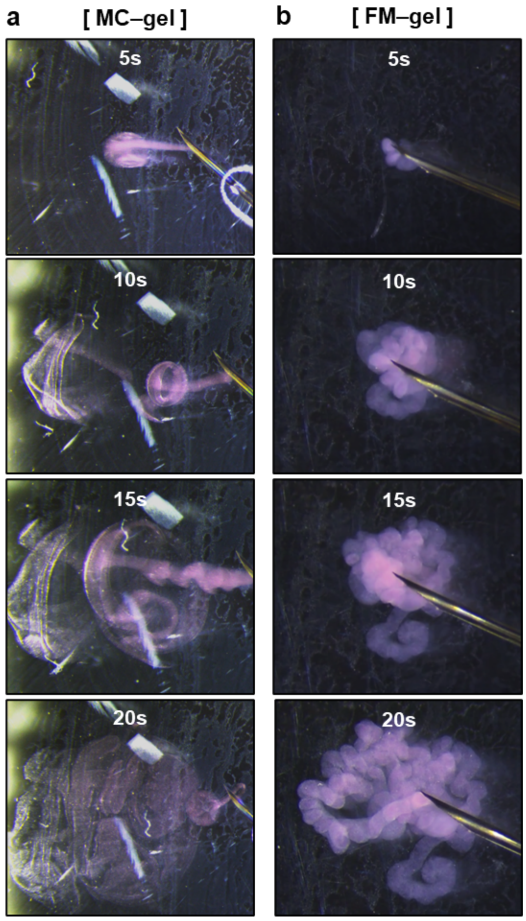


**Figure S1.** In situ gelation of hydrogel in PBS. The hydrogel containing DiI-loaded MCs (a) and FMs (b) was injected into a 37°C PBS using a syringe with 27G needle at a rate of 2 μl $s^{-1}$. The final concentration of multi-arm PEG linkers was 36 mg/mL, and the concentration of FM or MC solution was 30 mg/mL.

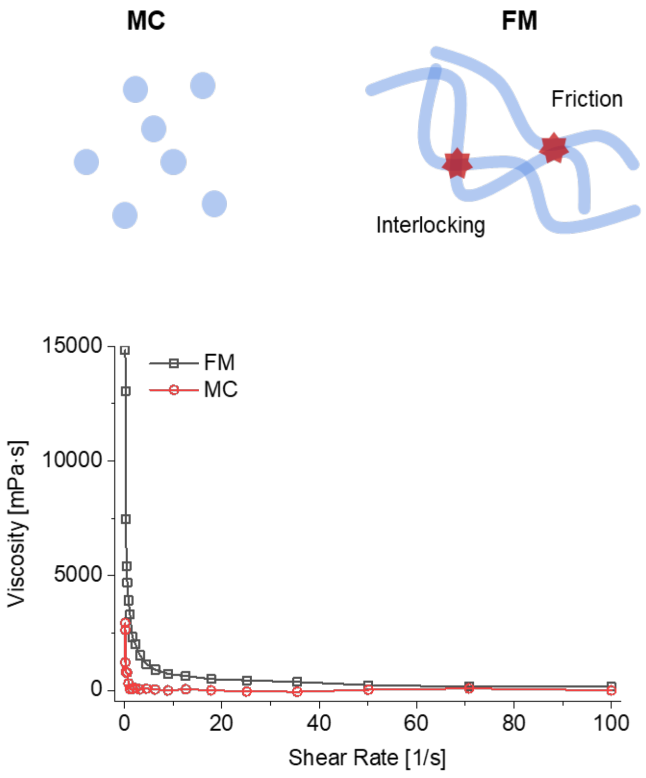


**Figure S2.** Viscosity of FM (black line) and MC (red line) solutions over a range of shear rates (0.1–100 $s^{-1}$). Both solutions are in a liquid state (not gels), and no PEG cross-linker is present. The block copolymer concentration of both nanocarrier solutions is 72.7 mg/ml.

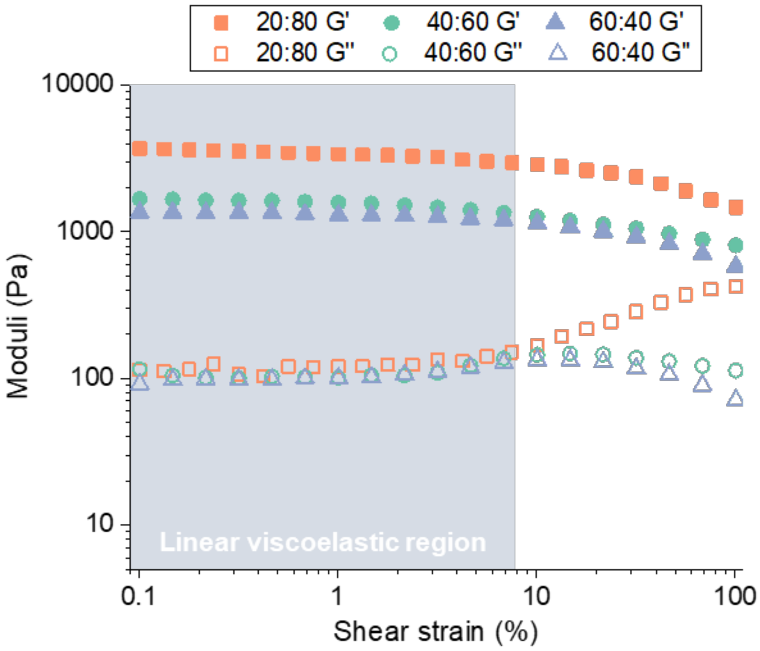


**Figure S3.** Strain-dependent oscillatory rheology of C-gels with different FM:PEG linkers volume ratios (Angular frequency = 6.28 rad s-1, 37°C).

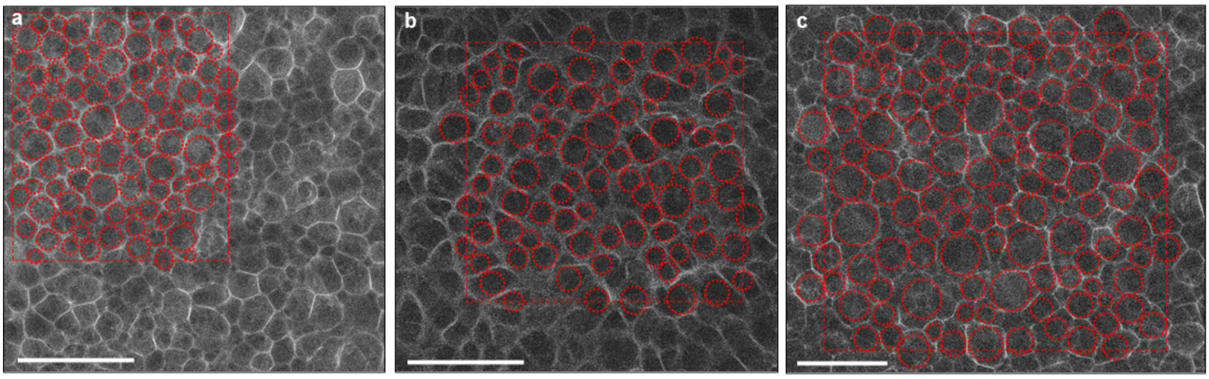


**Figure S4.** Examples of the evaluation of pore size. Pores used for determining the pore size using the ImageJ software are highlighted in the representative cryoSTEM images of C-gels with FM:PEG linkers ratios of 20:80 (a), 40:60 (b), and 60:40 (c) (v/v) as examples. The scale bar is 2 µm.

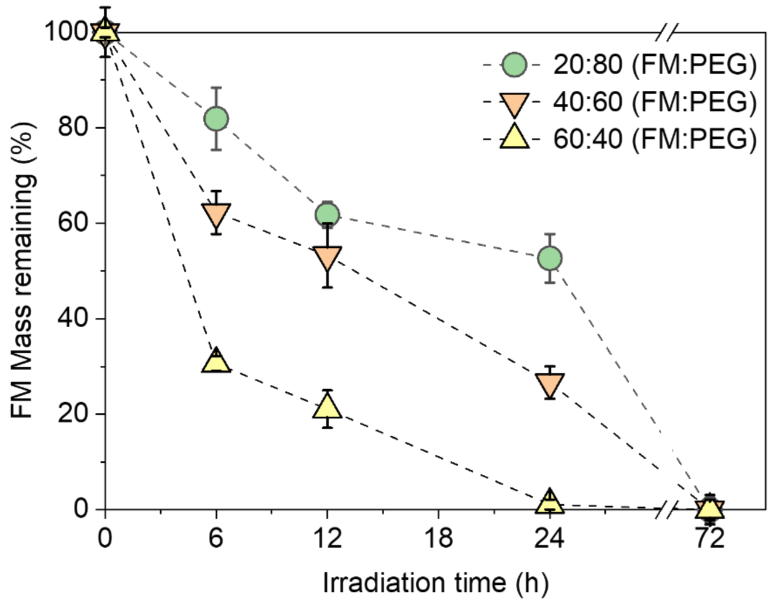


**Figure S5.** Percent mass remaining of FM in C-gels with FM:PEG linkers volume ratios of 20:80 (green), 40:60 (orange), and 60:40 (yellow) after light irradiation from 0 to 72 h (n = 3).

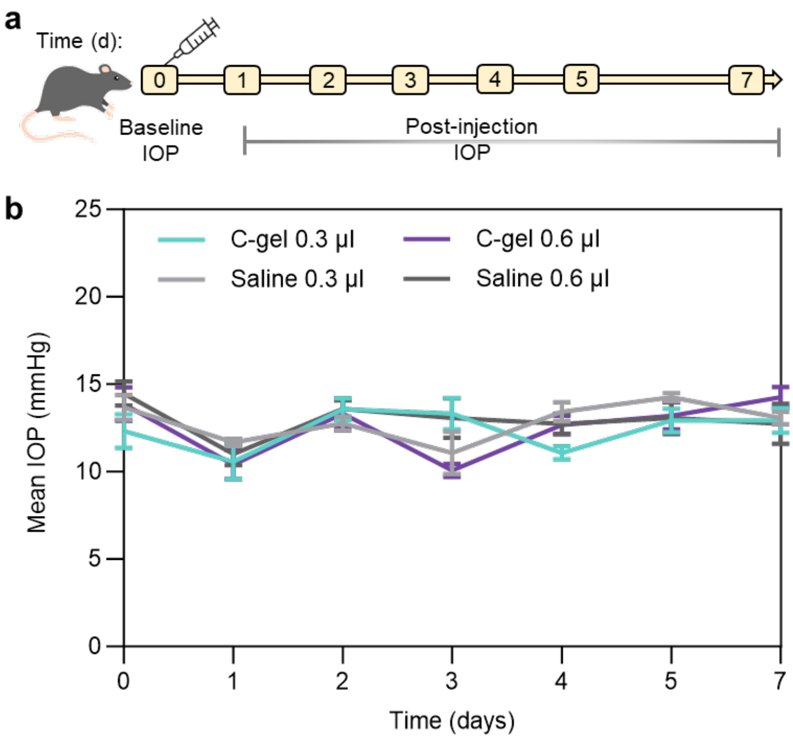


**Figure S6.** IOP response following intracameral injection of C-gel. (a) Schematic overview of a paired IOP study *in vivo*. 0.3 and 0.6 µl of C-gel were injected intracamerally, while an equal volume of balanced saline solution was injected into the contralateral eye. (b) Time course of IOP following intracameral injection. Injections were performed at day 0, and IOP was measured prior to injection, and assessed upto day 7, *n*=4 for 0.3 µl injected group and *n*=5 for 0.6 µl injected group. No statistically significant differences were observed between the two groups. Error bars represent s.e.m.

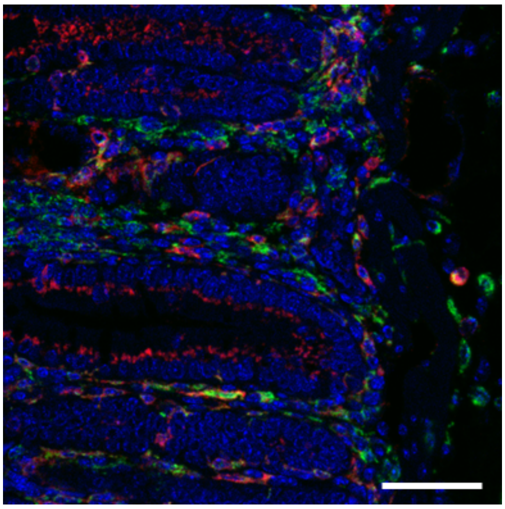

**Figure S7.** Positive control for CD68 and CD11c antibody staining. Representative immunofluorescence image of colon tissue from colitic mice used as positive control, confirming specific staining for CD68 (green) and CD11c (red). Nuclei were counterstained with DAPI (blue). Scale bar is 10 µm.
